# Analysis of cancer genomic amplifications identifies druggable collateral dependencies within the amplicon

**DOI:** 10.1101/2022.11.16.516737

**Authors:** G Pons, G Gallo-Oller, N Navarro, P Zarzosa, J Sansa-Girona, L García-Gilabert, A Magdaleno, MF Segura, J Sánchez de Toledo, S Gallego, L Moreno, J Roma

## Abstract

The identification of novel therapeutic targets for specific cancer molecular subtypes is crucial for the development of precision oncology. In the last years, CRISPR/Cas9 screens have accelerated the discovery and validation of new targets associated with different tumor types, mutations or fusions. However, there are still many cancer vulnerabilities associated with specific molecular features that remain to be explored. Here we used data from CRISPR/Cas9 screens in 954 cancer cell lines to identify gene dependencies associated with 16 common cancer genomic amplifications. We found that high-copy number genomic amplifications generate multiple collateral dependencies within the amplified region in 94% of cases. Further, to prioritize candidate targets for each chromosomal region amplified, we integrated gene dependency parameters with both druggability data and subcellular location. Finally, analysis of the relationship between gene expression and gene dependency leads to the identification of genes, the expression of which may constitute predictive biomarkers of dependency.

## Introduction

Focal chromosomal amplifications often drive an increase in the number of copies of certain oncogenes in malignant tumors^1^. As a consequence, cancer cells often become “addicted” to the overexpressed oncogenes increasing their malignancy and resistance to certain drugs^2^. Over the last 30 years, several studies have been conducted on assessing the effects of genomic amplifications in tumor prognosis. Overall, genomic amplifications such as *MYC, EGFR, CDK4* or *ERBB2* have been correlated with poorer survival in different tumors^3–6^. However, it should be noted that, regardless of whether genomic amplifications may be a factor of good or poor prognosis, only copy-number in a few amplified regions has been established as a biomarker for targeted therapies (e.g., *ERBB2* amplification and trastuzumab sensitivity)^7^. Nevertheless, at the same time as we quantify copy-number variation for molecular diagnosis in a wide variety of tumors, we still do not know the relevance in tumorigenesis of most genes embedded in the amplified regions. Currently, this hinders the clinical translationality of the molecular diagnosis towards the discovery and use of new targeted therapies for these patients. Thus, there is an increasing unmet necessity to identify specific druggable dependencies associated with chromosome amplifications recurrently detected in tumors. On this matter, CRISPR/Cas9 dropout screens have emerged as a useful approach to discover the role played by multiple proteins, as well as to identify new targets for tumors with specific molecular features^8–10^. However, some studies highlighted the generation of false-positive hits in CRISPR screens in cancer cell lines harboring genomic amplifications^11,12^. During the last few years, several methods such as Chronos algorithm have been developed to address some CRISPR screen artifacts simultaneously, including the nonspecific CRISPR-cutting induced toxicity observed in those sgRNAs targeting amplified regions^13,14^. Here we used CRISPR screen data corrected by Chronos algorithm to determine those genes on which cell lines harboring a high-copy number of a specific amplification are dependent. We also integrated gene dependency data with druggability information to especially prioritize those actionable targets for each chromosomal amplification (rationale overview in Fig. 1a). Finally, RNA-seq gene expression data was used to identify those genes in which mRNA levels may constitute a predictive biomarker of response to its inhibition. Overall, our analysis provides insights into the importance of a wide range of amplified genes with unknown function in cancer and identifies those selective gene dependencies associated with each chromosomal amplification to be further studied in the preclinical or clinical settings.

**Figure 1.**
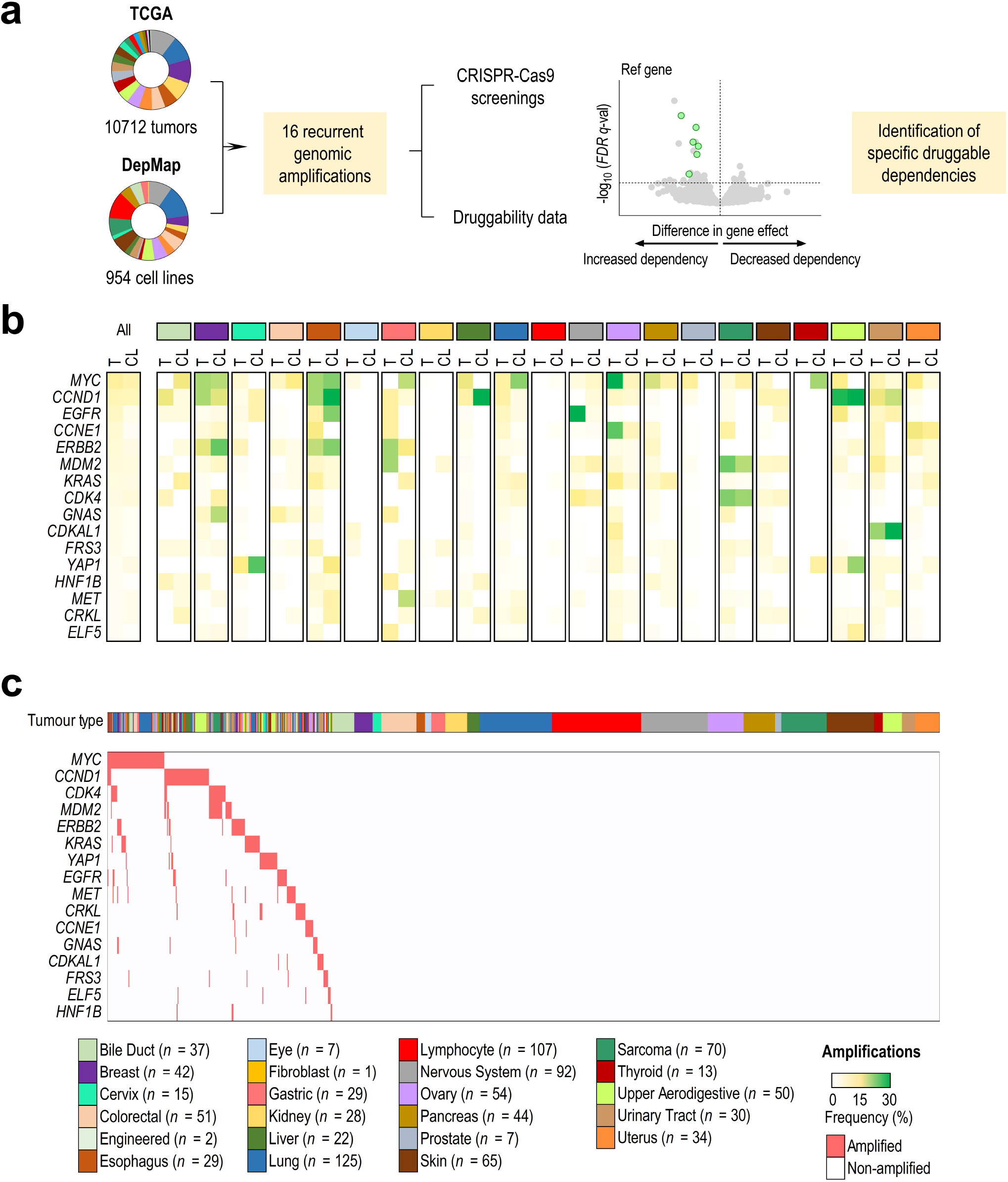
Cancer cell lines as a model to study tumor gene amplifications. (**a**) Conceptual overview of the approach used to identify specific drug vulnerabilities associated with 16 recurrent genomic amplifications. (**b**) Comparison of amplification frequencies between TCGA tumors (T) and DepMap cell lines (CL) from 21 lineages. A color gradient (from white to green) indicates the frequency of each amplification. (**c**) Amplification map of the DepMap cell lines (*n* = 954) used in the study. Cell lines were classified by the lineage of origin and ordered by the presence (red) or absence (white) of each amplification (≥ 6 copies).

## Materials and methods

### Selection of copy number amplifications in tumors and cell lines

Copy-number data from 10712 tumor samples were retrieved from a compilation of 32 TCGA Pan-cancer Atlas Studies (RRID:SCR_014555)^15,16^. Copy-number data for 954 cancer cell lines were retrieved from the DepMap dataset Copy Number 21Q4 Public^17^ (RRID:SCR_017655). Gene level copy-number data available in DepMap were relative to each cell line ploidy and log_2_-transformed with a pseudo count of 1 (log_2_ relative copy number + 1). To establish which cell lines harbored an amplification in each gene analyzed, absolute copy numbers (ACN) were calculated from relative copy numbers (RCN) using the following formula: ACN = cell line ploidy * ((2 ^ RCN) - 1). The following threshold was defined to consider a high-copy number amplification: RCN ≥ 2, which is equal to an ACN ≥ 6 copies in diploid cells. Since multiple genes are coamplified within a chromosomal amplification, a reference gene within the amplicon was used as a surrogate marker. Based on amplification frequency in tumors and cell lines, 16 reference genes were selected and used for the initial screening. For reference genes located in adjacent chromosomal bands (e.g., *CDK4* and *MDM2* located in chromosome 12q14 and 12q15, respectively), a degree of amplification co-occurrence was detected and taken into account in further analysis.

### Overall survival and disease-specific survival in tumor samples

Survival data from 10712 tumor samples were retrieved from TCGA Pan-cancer Atlas Studies (RRID:SCR_014555). Median overall survival (OS), disease-specific survival (DSS) and hazard ratios (HR) were calculated to assess the effect of the 16 amplifications in patient prognosis. Samples bearing amplifications in the reference genes of each region were compared to samples with none of the amplifications. Stratification was applied to better understand the significance of each amplification within each tumor type. Significance between survival probability curves was determined using the log-rank test.

### Comparison of similarity in genomic amplification between tumors and cell lines

To compare amplifications between tumors and cell lines, TCGA tumors (*n* = 10712) and DepMap cell lines (*n* = 954) were grouped into 25 and 23 lineages respectively, 21 of which were shared among them (Supplementary Tables S1A and S1B). Subsequently, the amplification frequencies in tumors and cell lines for each reference gene and lineage were correlated. A greater correlation between amplification frequencies in tumors compared to cell lines meant a greater similarity among them in terms of gene amplification. Therefore, in relation to the calculated Pearson’s correlation coefficient (*r*) three thresholds were established: high (*r* ≥ |0.7|), moderate (|0.7| < *r* ≥ |0.4|) and weak or negligible correlation (*r* < |0.4|).

### Identification of coamplified genes within a region

To elucidate the relevance of each gene amplified within the regions analyzed in cell survival, it was necessary to identify which genes were coamplified with each reference gene. To address that, we correlated copy-numbers of the reference genes with copy-numbers of the remaining genes in cell lines. Genes whose copy-numbers showed an *r* ≥ 0.7 with the reference gene copy-numbers were considered coamplified genes (Supplementary Table S2). We also integrated data regarding chromosome location and gene type by using BioMart^18^ (RRID:SCR_002987) and Genecards^19,20^ (RRID:SCR_002773).

### Screening of gene dependencies associated with gene amplifications

To analyze those dependencies associated with each gene amplification, gene effect data of 17385 genes in 954 cell lines were retrieved from the DepMap dataset CRISPR 21Q4 Public+Score, Chronos^13,21,22^ (RRID:SCR_017655). Subsequently, the differences in gene effects between those cell lines harboring each of the 16 amplifications (ACN ≥ 6 copies) with respect to those cell lines that did not harbor each of them (ACN < 6 copies) was calculated. The significance between gene effects was determined using a two-tailed Student’s t-test followed by Benjamini-Hochberg correction to obtain FDR *q*-values. Then, for each amplification, we performed an overlap analysis between those significant dependent genes (query genes, (k)) and C1 positional gene sets (gene set, (K)) using MSigDB^23^ (RRID:SCR_016863). C1 positional gene sets are a compendium of 299 gene sets accounting for those human genes annotated on the GCh38.p13 reference chromosome bands. Thus, overlap analysis was performed to find chromosomal localization overlaps within those significant dependent genes. The ratio between the number of query genes (k) overlapped with the number of genes within each gene set (K) was plotted on the x-axis whereas the gene set name was plotted on the y-axis. FDR *q*-value was indicated by a color gradient and gene set size by a size dot gradient. Graphpad Prism 6.01 (RRID:SCR_017655) was used to generate the volcano plots and overlap plots. In addition, to confirm an enrichment of coamplified genes in the gene dependencies obtained, for each amplified region we ran a preranked GSEA^24^ (RRID:SCR_003199), comparing the preranked list of dependent genes (ordered by *q*-value) with the corresponding gene set of coamplified genes. The following parameters were set: 10000 permutations, weighted enrichment statistic, and meandiv as normalization mode. Finally, we used the normalized enrichment score (NES) and the associated FDR *q*-value to statistically interpret the enrichment plots obtained.

### Prioritization of candidate targets for each chromosome amplification

To prioritize candidate targets for each chromosome amplification, we integrated druggability and subcellular localization data from canSAR^25^ (RRID:SCR_006794) for the 923 significant gene dependencies identified (corresponding to 770 different genes) (Supplementary Table S3). Considering that cellular localization is important for target druggability, proteins secreted or located in the cell membrane were selected from those located in other cellular compartments. Then, structure-based ligandability scores, ranked from low-ligandability (−3) to high-ligandability (3), were added using a color-gradient. To verify the results obtained in the initial pan-cancer screening, we also compared the difference in gene effects of those prioritized genes between amplified and non-amplified cell lines in selected lineages. In the case of *SLC26A10* and *CALM1P2*, as copy number data was not available, we used as a surrogate marker of amplification *B4GALNT1* and *EGFR* respectively, which are genes closely located to both genes of interest. The significance between gene effects was determined using a two-tailed Student’s t-test. Graphpad Prism 6.01 (RRID:SCR_017655) was used to generate the plots.

### Correlation between gene dependencies and gene expression

RNA-seq expression data of prioritized genes were retrieved from the DepMap dataset Expression 21Q4 Public^17^ (RRID:SCR_017655). Gene expression data were log_2_ transformed with a pseudo count of 1 (log_2_ TPM+1). Since the immediate downstream effect of amplification is overexpression, we first determined which prioritized genes were overexpressed when amplified. To address that, for each prioritized gene, relative copy numbers were correlated with gene expression in selected tumor types. In the case of SLC26A10 we could not perform any correlation as neither copy number nor gene expression was available. In the case of *FKBP9* and *CALM1*, we correlated their expressions with *FKBP9P1* and *CALM1P2* copy numbers, respectively, as *FKBP9P1* and *CALM1P2* were found to be amplified but not *FKBP9* and *CALM1*. Then, to determine whether mRNA levels could be a predictive biomarker of gene dependency, gene expression was correlated with gene effect data in specific tumor categories because of putative lineage-specific differences in gene expression. The *p*-values obtained were corrected using Benjamini-Hochberg correction to obtain FDR *q*-values. Graphpad Prism 6.01 (RRID:SCR_017655) was used to generate the plots.

## Data availability statement

The data generated in this study are available within the article and its supplementary data. All gene sets generated regarding the coamplified genes within each chromosomal region will be submitted for its inclusion in MSigDB (RRID:SCR_016863), immediately after the acceptance of the manuscript. The data analyzed from TCGA (RRID:SCR_014555), DepMap (RRID:SCR_017655), CanSAR (RRID:SCR_006794), BioMart (RRID:SCR_002987) and Genecards (RRID:SCR_002773) were publicly available from their public websites as stated in the corresponding section in Materials and Methods.

## Results

### Cancer cell lines as a model to study tumor gene amplifications

Before gene dependency analysis, an initial key question was whether gene amplifications detected in cell lines reflected those observed in the tumors from which they were derived, or whether particular amplifications have been selected in cell culture implying an artefactual genetic bias. Correlations between gene amplification frequencies in tumors and cell lines showed that gene amplification profiles in cell lines resembled those observed in same lineage tumors (Fig. 1b). In particular, we found that amplification frequencies between tumors and cell lines showed a high correlation in 7/17 (41%) lineages, a moderate correlation in 6/17 (35%) lineages and a weak correlation in 4/17 (24%) lineages. It was not possible to correlate gene amplification frequencies in ocular, kidney, lymphoid and prostate lineages since in these pathologies gene amplifications were found to be a rare phenomenon. To clearly visualize the data analyzed, a gene amplification map was generated for cell lines (Fig. 1c) and tumors (Supplementary Fig. S1a). At this point, we also aimed to study the relevance of each amplification in tumor survival comparing those tumors that harbor each gene amplification with those that did not harbor any. In the pan-cancer analysis, a decreased probability of overall survival (OS) and disease-specific survival (DSS) was found in gene amplified tumors in comparison with non-amplified ones (Supplementary Fig. S2a – b), being *EGFR, CDK4, MDM2, KRAS* and *CCNE1*-amplified tumors those associated with a poorer prognosis (Supplementary Fig. S2c – d). Survival analysis by tumor type revealed that *EGFR* amplifications in low-grade glioma and head and neck cancers, *CCNE1* amplifications in ovarian and uterine cancers, and *CDK4* amplifications in sarcomas were associated with a low probability of OS suggesting that only certain amplifications in particular tumor types are relevant in prognosis (Supplementary Fig. S2e – t).

### Chromosome amplifications generate collateral dependencies within the amplicon

To elucidate, within the regions analyzed, the importance of each amplified gene in cell survival, all coamplified genes with the reference gene were identified. For each region amplified, we found a distinct number and frequency of protein-coding genes coamplified (Supplementary Fig. S3a). The gene type (protein-coding genes, RNA genes or pseudogenes) was considered, as for some non-coding genes copy-number data were available but not CRISPR-Cas9 gene effect data. For this reason, we could not obtain relevant data for *MYC* amplicon (chr8p24), since *MYC* and *POUF5F1B* were the only protein-coding genes coamplified in this region (Supplementary Fig. S3b-c). In the remaining 15 regions, we compared the differences between CRISPR-Cas9 gene effects in those cells with the reference gene amplified with respect to the non-amplified ones. Thus, we identified those genes whose inhibition would impair cell survival of cancer cells bearing each of the amplifications analyzed (Fig. 2a-i and Supplementary Fig. S3d-i). In addition, we used MSigDB to find overlapped chromosome localizations within those gene dependencies that were significant in the analysis (*q* < 0.05). Interestingly, most dependent genes were located within the same chromosomal band or, in some cases, in adjacent chromosomal regions (Fig. 2a-i). A preranked GSEA was also performed to confirm an enrichment of coamplified genes among the most significant gene dependencies for each region analyzed (Supplementary Fig. S4a-i). Of note, not all coamplified genes showed a collateral gene dependency, thus demonstrating the importance of finding those particular gene amplifications with greater relevance in cell survival. In particular, *CDK4*-amplified cell lines were highly dependent on *SLC26A10, TSPAN31* and *CPM*, all of them within 12q13-q15 (Fig. 2a). *MDM2*-amplified cell lines, which showed a high co-occurrence with *CDK4* amplification, showed a high dependency towards the same genes as CDK4-amplified cell lines and, in this case, also to *BEST3* (Supplementary Fig. S3d). *KRAS*-amplified cell lines were especially dependent on *CASC1* (Fig. 2b), and *EGFR*-amplified cell lines were highly dependent on *FKBP9* and *CALM1*. Intriguingly, neither *FKBP9* nor *CALM1* were located near or within the 7p11 region, but two of their pseudogenes, *FKBP9P1* and *CALM1P2*, were found to be coamplified with *EGFR* (Fig 2c). *CCND1*-amplified cell lines were especially vulnerable to *CCND1* depletion, but also to *FGF19* and *FGF4* depletion, while *ERBB2*-amplified and *CCNE1*-amplified cell lines strongly depend on *ERBB2* and *CCNE1*, respectively (Fig. 2d-f). In the case of *FRS3*-amplified cell lines, a significant sensitivity to *PRICKLE4, FOXP4* and *FRS3* depletion was observed, whereas *GNAS*-amplified ones strongly depend on *TFAP2C;* and *YAP1*-amplified cell lines depend on *YAP1* or *MMP27* (Fig. 2g-i). Additional relevant gene dependencies associated with amplifications in *CDKAL1, ELF5, CRKL, MET* or *HNF1B* are shown in Supplementary Fig. S3e-i.

**Figure 2.**
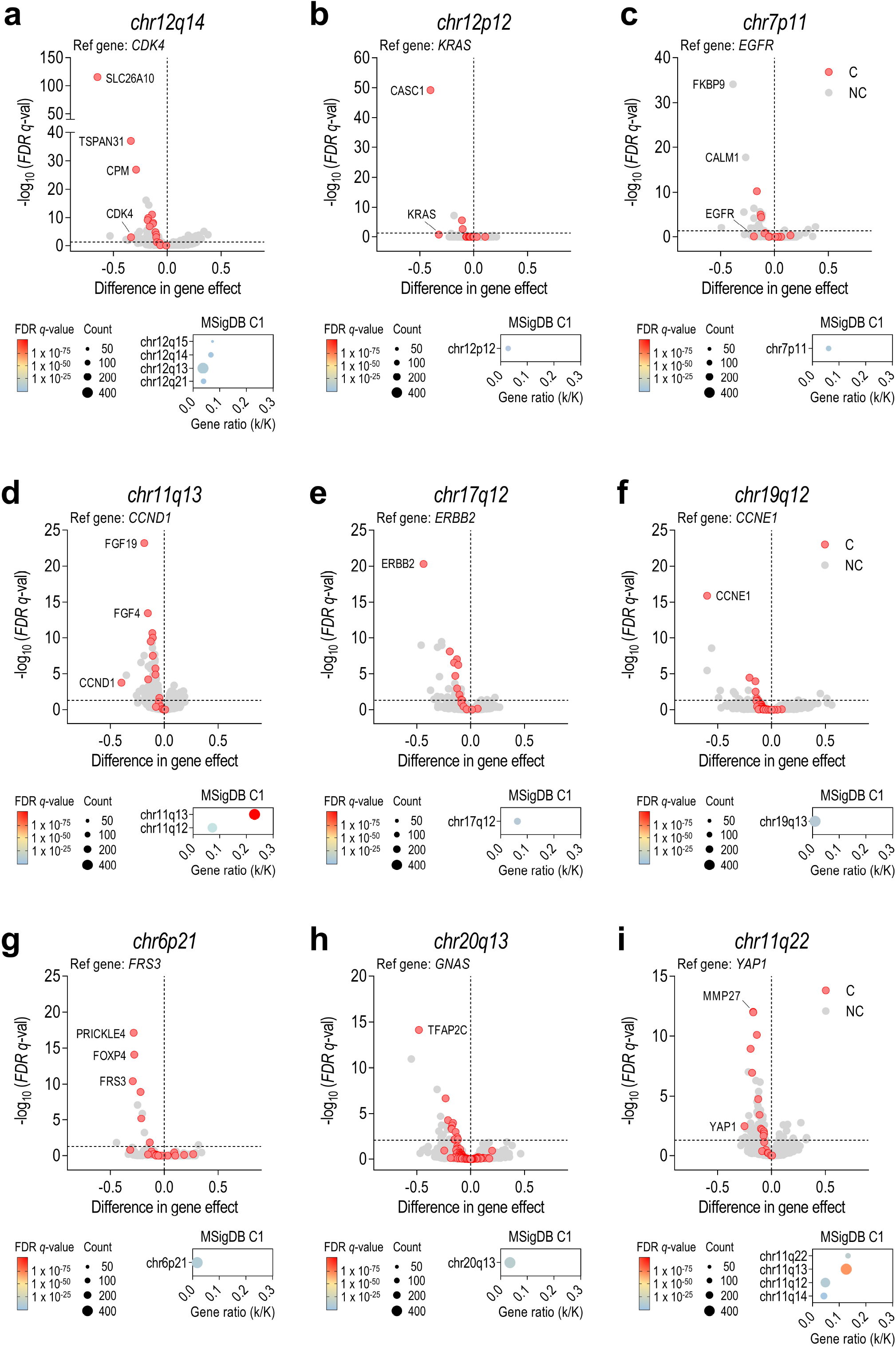
Identification of amplification-associated dependencies reveal the importance of coamplified genes. Results are shown for the following amplifications: (**a**) *CDK4*, chr12q14 (**b**) *KRAS*, chr12p12 (**c**) *EGFR*, chr7p11 (**d**) *CCND1*, chr11q13 (**e**) *ERBB2*, chr17q11 (**f**) *CCNE1*, chr19q12 (**g**) *FRS3*, chr6p21 (**h**) *GNAS*, chr20q13 and (**i**) *YAP1*, chr11q22. **Top**: Volcano plots showing the difference in CRISPR-Cas9 gene effect between cell lines harboring amplifications (≥ 6 copies) or not (< 6 copies). Genes were classified in coamplified (C, red dots) and non-coamplified genes (NC, grey dots). Statistical significance (*q* < 0.05) was determined using two-tailed t-tests followed by Benjamini-Hochberg correction to obtain FDR *q*-values. **Bottom**: MSigDB overlap plots revealed an enrichment of amplification-associated dependencies in genes located within the same chromosomal band or in adjacent chromosomal regions. Gene ratio (k/K) is referred to the number of query genes (k) overlapped with the number of genes within each gene set (K). FDR *q*-value is indicated by a color gradient and gene set size by a size dot gradient.

### Some collateral dependencies generated by amplification are druggable

Once gene dependencies associated with each gene amplification were characterized, we wondered which ones might be prioritized as putative new targets. Prioritization for the significant gene dependencies previously found was based on both the subcellular localization and druggability score of each particular target (Fig. 3a-b). Dependent genes were classified by the subcellular localization of the encoded protein, considering that membrane and secreted proteins are more accessible to be inhibited by small molecules or antibodies. Druggability scores were particularly taken into consideration in those dependent genes codifying proteins located in the cytoplasm, nucleus or organelles. Among dependent genes codifying proteins located in the membrane, we highlighted, as priority targets, the following genes: *SLC26A10, CPM, TSPAN31* and *BEST3* (12q13-15), *FGF19* and *FGF4* (11q13), *ERBB2* (17q12), *FRS3* (6p21), *PAMR1* (11p13) and *MET* (7q31) (Fig. 3a-b,

**Figure 3.**
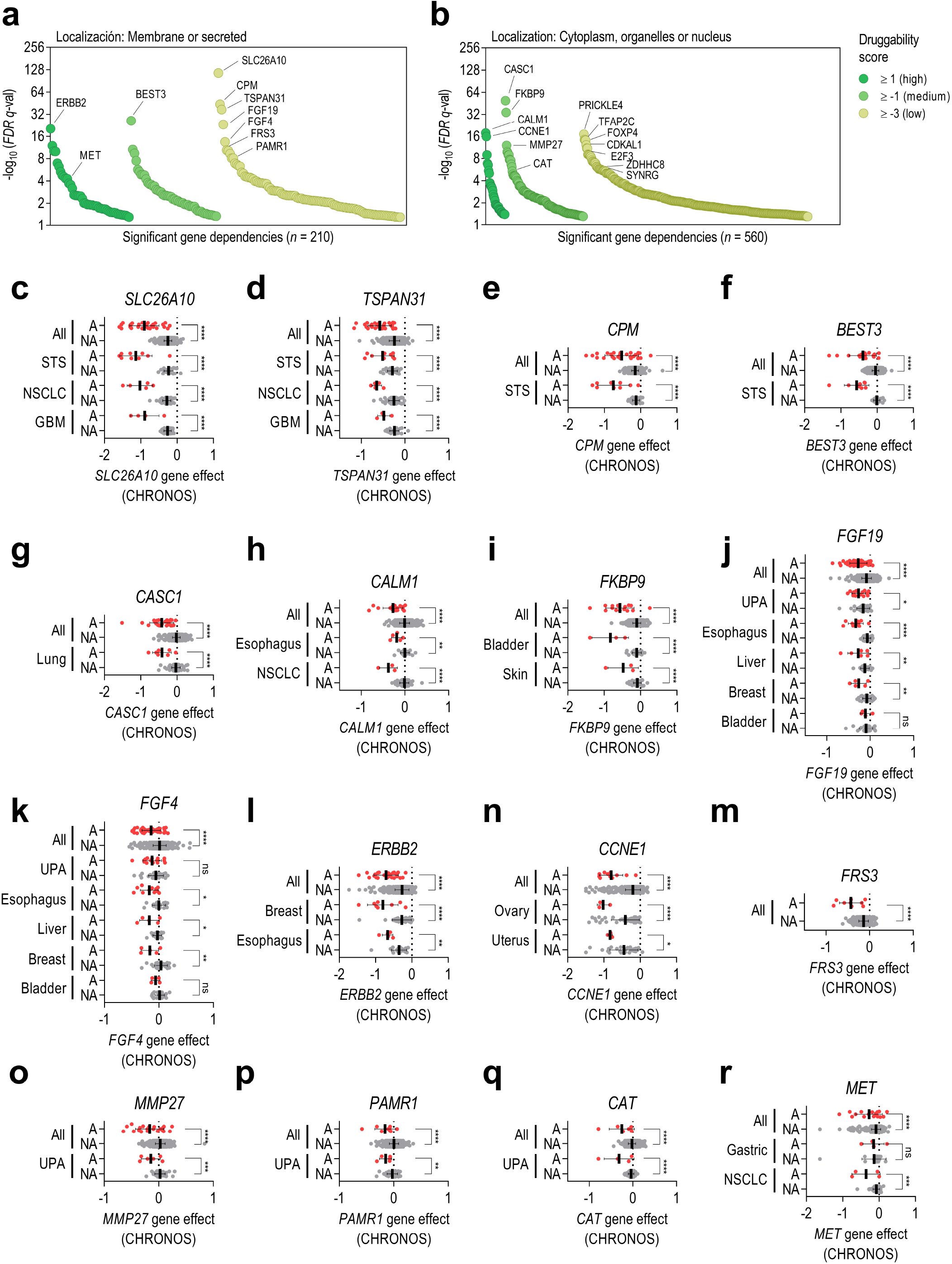
Prioritization of candidate targets for each chromosome amplification. CanSAR druggability scores (from +3 to -3) were integrated for all significant dependent genes previously identified. A higher druggability score indicates a higher probability of the protein codified to harbor ligandable pockets within its 3D structure. Dependent genes were classified in (**a**) genes codifying for membrane or secreted proteins or (**b**) genes codifying for proteins located in cytoplasm, organelles or nucleus. (**c-r**) Difference in gene effects of the prioritized genes between amplified (A, red dots) and non-amplified (NA, grey dots) cell lines from specific tumor types. Results are shown for the following genes: (**c**) *SLC26A10*, (**d**) *TSPAN31*, (**e**) *CPM*, (**f**) *BEST3*, (**g**) *CASC1*, (**h**) *CALM1*, (**i**) *FKBP9*, (**j**) *FGF19*, (**k**) *FGF4*, (**l**) *ERBB2*, (**m**) *CCNE1*, (**n**) *FRS3*, (**o**) *MMP27*, (**p**) *PAMR1*, (**q**) *CAT*, and (**r**) *MET*. Statistical significance (p<0,05) was determined using two-tailed t-tests. STS: soft-tissue sarcoma; NSCLC: non-small cell lung cancer; GBM: glioblastoma; UPA: upper aerodigestive tumors.

Supplementary Table S3). Regarding dependent genes encoding proteins not located in the membrane nor secreted, we prioritized *CASC1* (12p12), *CALM1* and *FKBP9* (*FKBP9P1* and *CALM1P2* pseudogenes located in 7p11), *CCNE1* (19q12), *MMP27* (11q12), and *CAT* (11p13) (Fig. 3a-b, Supplementary Table S3). To confirm the results obtained in the initial pan-cancer screening, we also compared the difference in gene effects of those prioritized genes between amplified and non-amplified cell lines in specific tumor types (Fig. 3c-r). Differences in gene effects between amplified versus non-amplified cell lines in distinct tumor types were similar, as it was observed in *SLC26A10, TSPAN31, CALM1, FKBP9, ERBB2* or *CCNE1*, suggesting that the biological consequences derived from the knockout of these prioritized genes are, in general, independent of lineage. However, in some tumor types no differences in gene effect were observed as exemplified by *FGF19* and *FGF4* in bladder tumors or *MET* in gastric tumors.

### mRNA gene expression levels correlate with gene dependency in some prioritized genes

Since the immediate downstream effect of gene amplification is gene overexpression, we aimed to determine whether gene expression levels could be a predictive biomarker of gene dependency. To address that, we first determined which of the prioritized amplified genes were overexpressed when amplified in specific tumor categories. For each prioritized gene we correlated its relative expression with its relative copy number (Supplementary Fig. S5a-o). Then, relative expression was correlated with gene dependency data in cell lines from selected lineages (Fig. 4A-O). Overall, the strongest correlations between expression and dependency were found in those prioritized genes whose copy number strongly correlated with gene expression (*BEST3, CPM, ERBB2, CCNE1* or *TSPAN31*) (Fig. 4a-e and Supplementary Fig. S5a-e. However, in some cases, such as *CALM1, FGF4, FKBP9, PAMR1* or MMP27, there was no clear correlation between gene amplification and gene overexpression nor between gene expression and gene dependence (Fig. 4k-o and Supplementary Fig. S5k-o). The fact that an increase in copy number does not always cause an increase in gene expression may be due to existing negative-feedback mechanisms to control gene expression at transcript level for some genes. Our results highlight, at least in the prioritized genes analyzed, the importance of taking into account gene copy number as a predictive biomarker of gene dependence instead of only consider gene expression.

**Figure 4.**
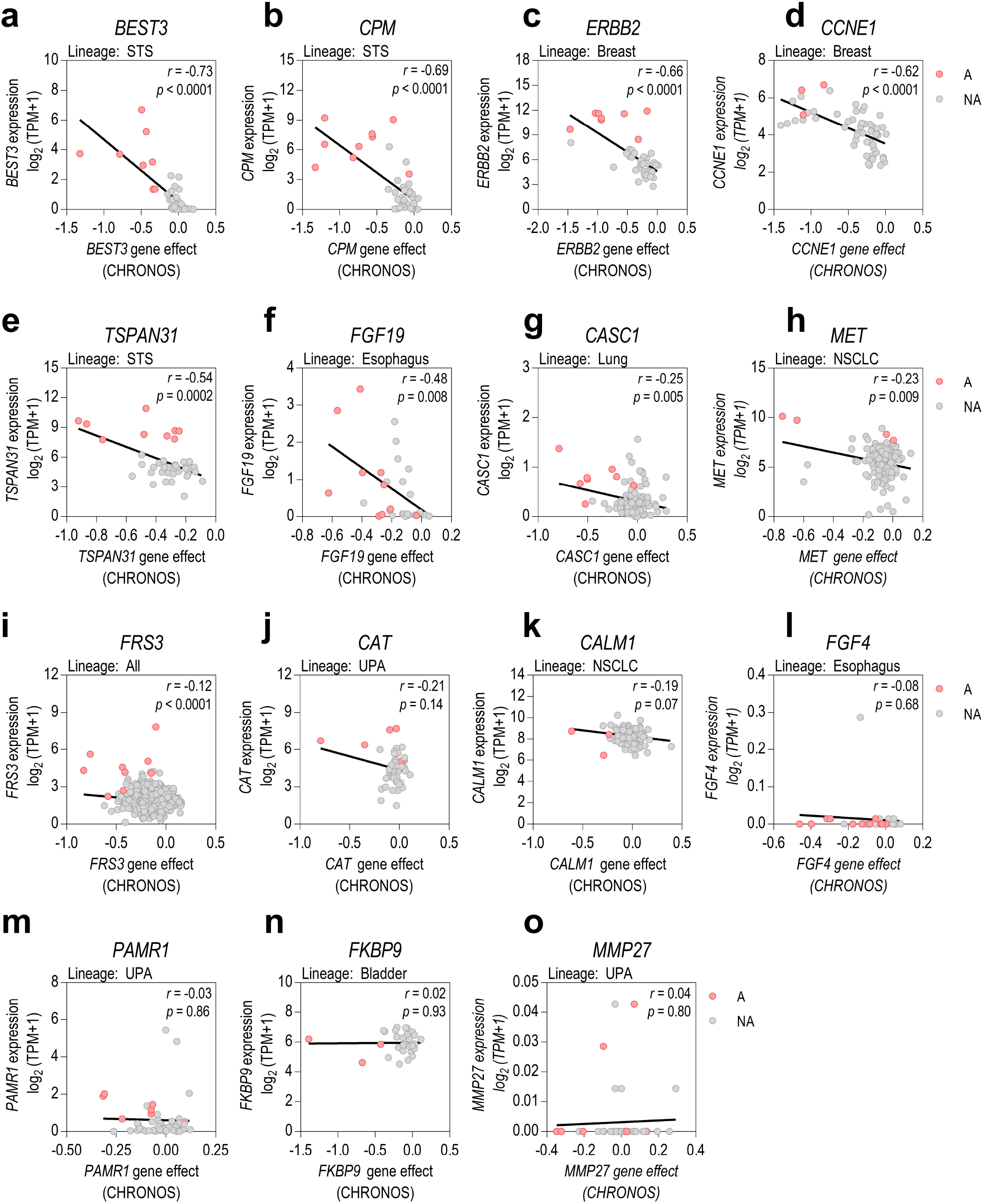
Assessment of gene expression as a predictive biomarker of gene dependency. (**a-o**) Pearson’s correlations between relative gene expression and gene knockout effects are shown for the prioritized genes in cell lines from selected lineages. Cell lines were also classified in amplified (A, red dots) and non-amplified genes (NA, grey dots). STS: soft-tissue sarcoma; NSCLC: non-small cell lung cancer; UPA: upper aerodigestive tumors.

## Discussion

In this study, we used data from CRISPR-Cas9 screens in 954 cancer cell lines to identify gene dependencies associated with 16 common genomic amplifications detected in tumors. We observed a greater dependence among coamplified genes within each of the regions analyzed with respect to non-coamplified genes, suggesting that particular coamplified genes with previously unknown function in tumor malignancy confer a higher survival capacity to cancer cells harboring each amplification. Although some drugs have already been developed for some previously known amplification-associated dependencies, such as ERBB2 or MET, there are no specific drugs for inhibiting most of the gene dependencies described herein. In this regard, we integrated druggability data and subcellular localization to better select those gene dependencies that might be prioritized for further research and drug development. Our analysis raises the possibility to specifically target many new vulnerabilities associated with recurrent chromosome amplifications detected in tumors and supports previous results research on targeting particular associated dependencies. Additionally, we also determined for which dependent genes mRNA levels could constitute a predictive biomarker of their dependency, either independently or as a complement of copy-number levels. We believe that these results could be reproduced and extended to almost all chromosomal amplifications detected in tumors, but a representative subset of cell lines harboring those amplifications would be required. Moreover, it would also be relevant to analyze the effects derived from depleting ncRNA genes, especially in amplified regions presenting low frequency of protein-coding genes, to obtain additional new targets. Overall, the discovery of new vulnerabilities associated with recurrent amplifications detected in tumors might entail a major advance in the development of new therapies against cancer, thus contributing to the progress of precision medicine.

## Acknowledgments

We specially wish to thank the Cancer Dependency Map Project team for making all the data generated available to the scientific community. Without their previous efforts to integrate CRISPR-Cas9 loss of function screenings with multi-omics data in a wide variety of cancer cell lines, this work would not have been accomplished. We also thank Ms. Helena Kruyer for her help with the English version of this manuscript.

## Author’s contributions

Conceptualization: J.R. and G.P.; methodology: G.P., G.G-O and A.M.; formal analysis: G.P., G.G-O., N.N., P.Z. and J.S-G; data curation: G.P., G.G-O., N.N., P.Z., J.S-G and L.G-G; writing— original draft preparation: G.P., G.G-O. and J.R.; writing—review and editing: G.P., G.G-O., J.R., L.M., MF.S., and S.G.; supervision: J.R. and L.M.; project administration: J.R.; funding acquisition: J.R., L.M., J.SdT. and S.G. All authors have read and agreed to the published version of the manuscript.

## Figure Legends

**Supplementary Figure S1.**
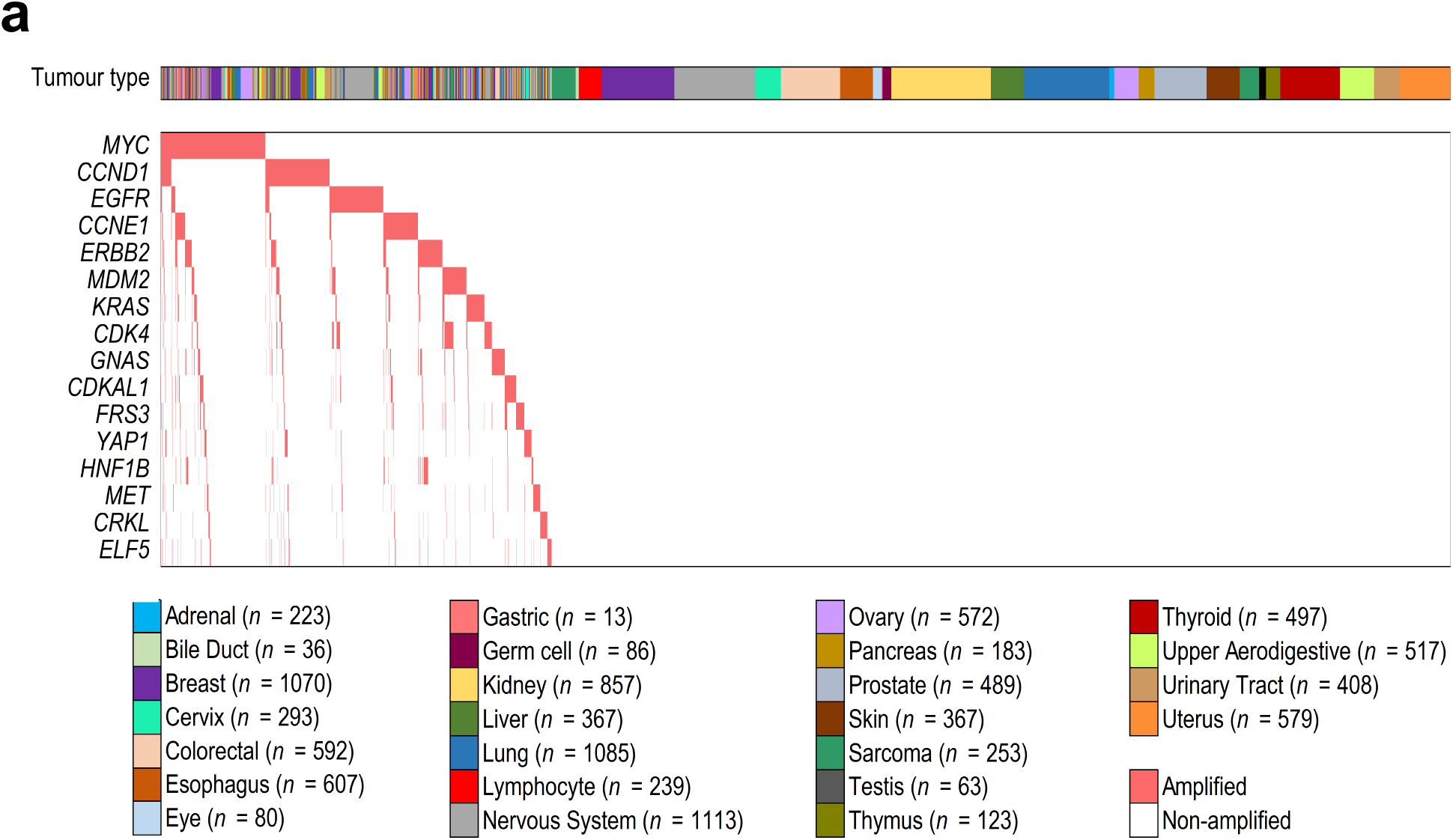
Genomic amplifications in TCGA tumors. (**a**) Amplification map of the TCGA tumors (*n* = 10712) used in the study. Tumors were classified by the lineage of origin and ordered by the presence (red) or absence (white) of each amplification.

**Supplementary Figure S2.**
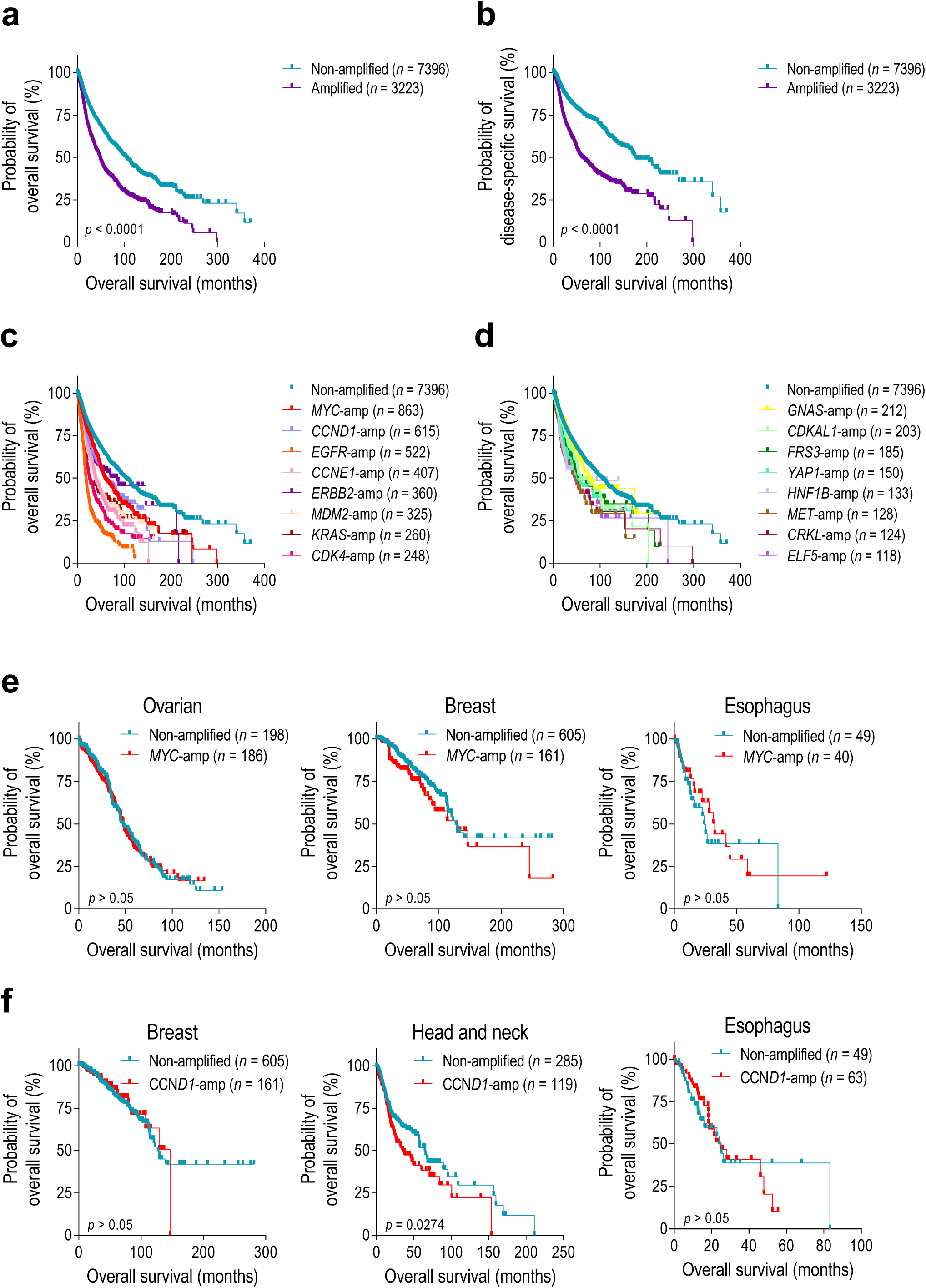
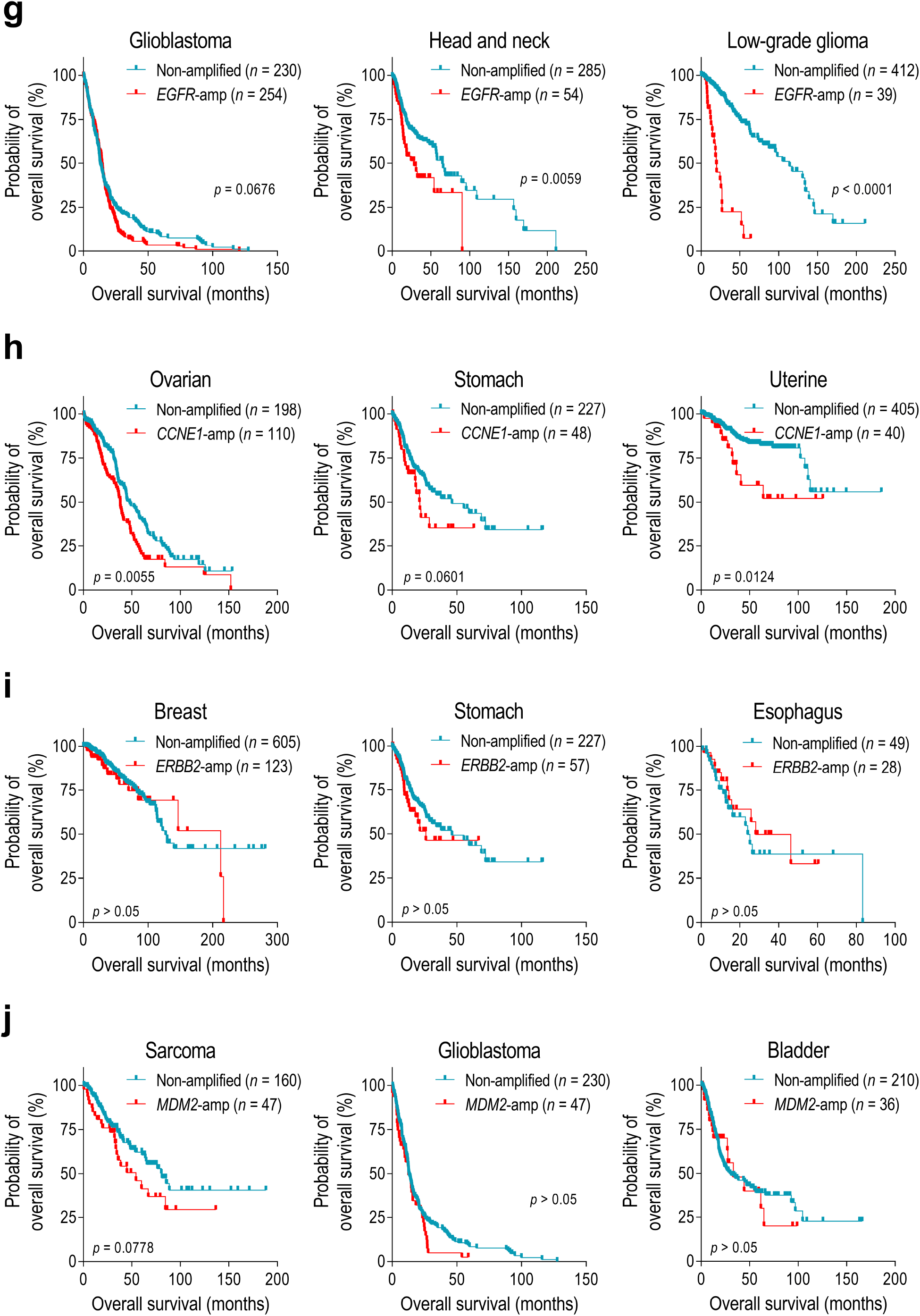
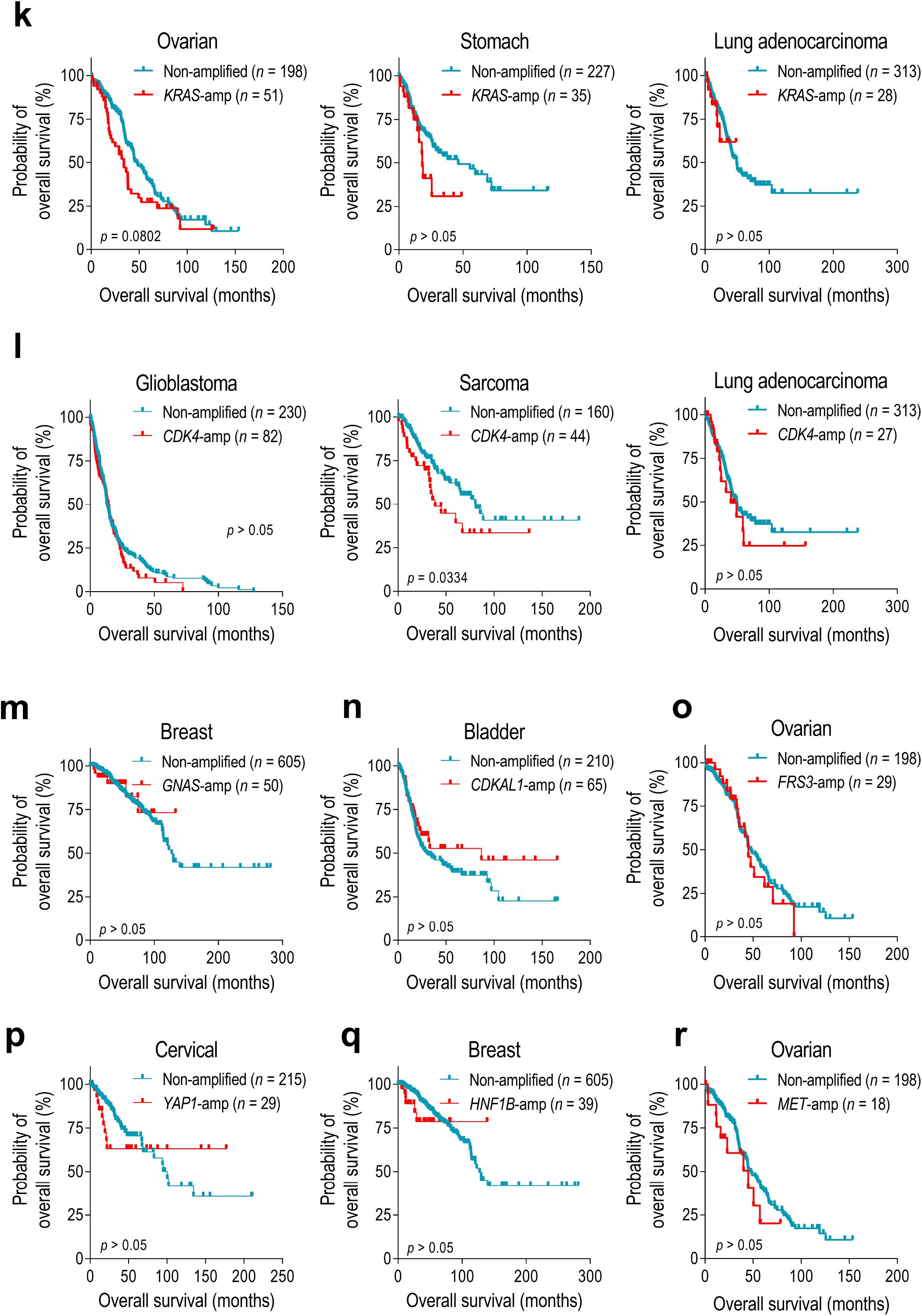
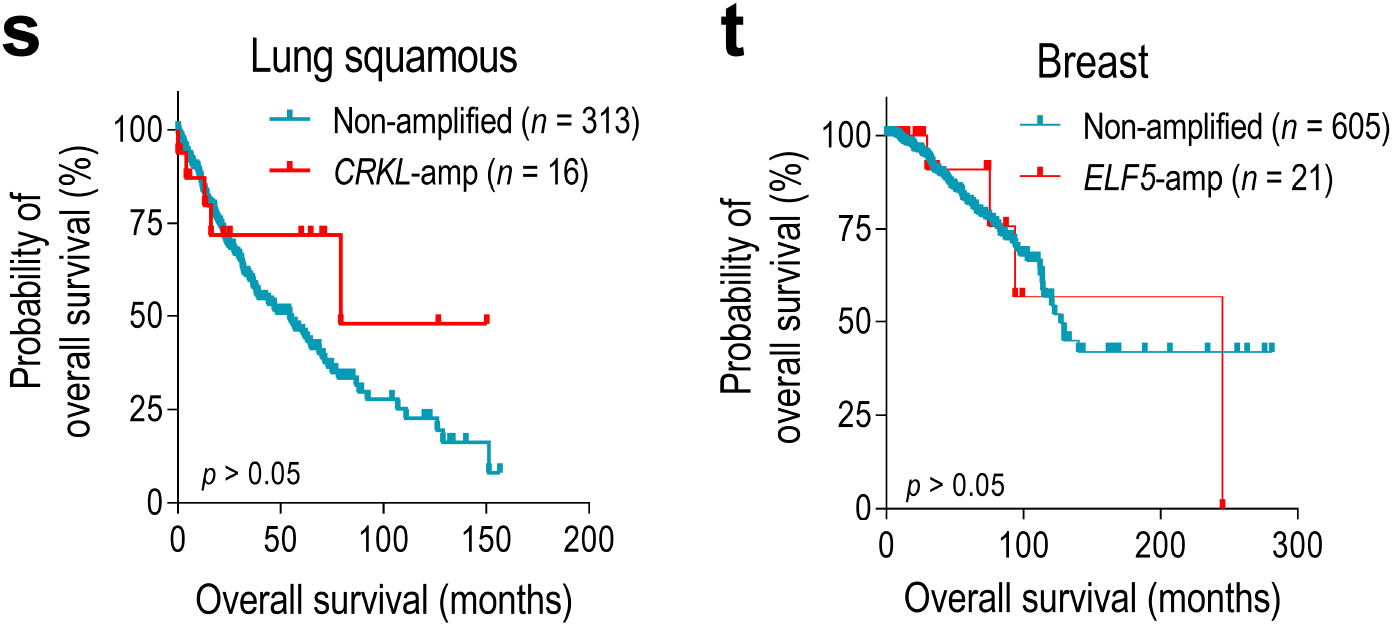
Survival associated with gene amplifications in TCGA tumors. (**a-b**) Kaplan-Meier curves showing the probability of patient’s overall survival and disease-specific survival comparing those tumors which harbor any of the gene amplifications analyzed with those which did not harbor any of them. (**c-d**) Kaplan-Meier curves showing the probability of patient’s overall survival comparing those tumors which harbor each gene amplification with those which did not harbor any of them. (**e-t**) Kaplan-Meier curves showing the probability of patient’s overall survival comparing those tumors which harbor each gene amplification with those which did not harbor any of them in selected lineages where the amplification was frequently detected. Considering the number of patient samples harboring each amplification, 3 tumor types were selected regarding amplifications more frequently detected in TCGA tumors and 1 tumor type in amplifications less frequently detected. Statistical significance (*p* < 0.05) was determined using log-rank test.

**Supplementary Figure S3.**
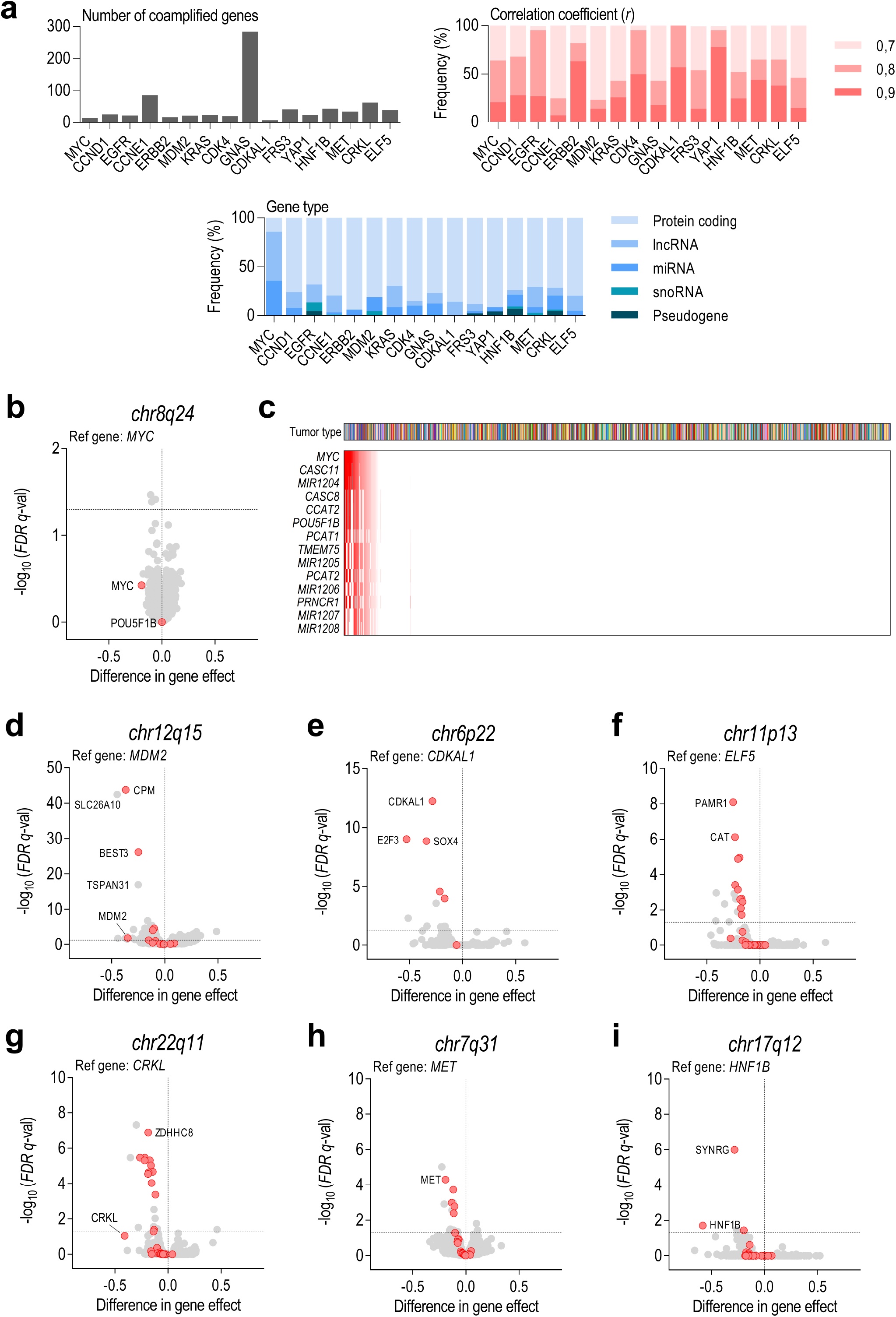
Identification of amplification-associated dependencies reveal the importance of coamplified genes. (**a**) Left: Number of coamplified genes with the reference gene. Right: Frequency of coamplified genes by its correlation coefficient with the reference gene. Below: Frequency of coamplified genes by its gene type (protein coding, lncRNA, miRNA, snoRNA or pseudogenes). (**b**) Volcano plots showing the difference in gene effect between cell lines harboring *MYC* amplifications or not. (**c**) The majority of coamplified genes with *MYC* (*r* > 0,7) are non-coding genes. (**d-i**) Volcano plots showing the difference in gene effect between cell lines harboring the remaining 5 amplifications analyzed: (**d**) *MDM2*, chr12q15 (**e**) *CDKAL1*, chr6p22 (**f**) *ELF5*, chr11p13 (**g**) *CRKL*, chr22q11 (**h**) *MET*, chr7q31 (**i**) *HNF1B*, chr17q12. Statistical significance (*q* < 0.05) was determined using two-tailed t-tests followed by Benjamini-Hochberg correction to obtain FDR *q*-values.

**Supplementary Figure S4.**
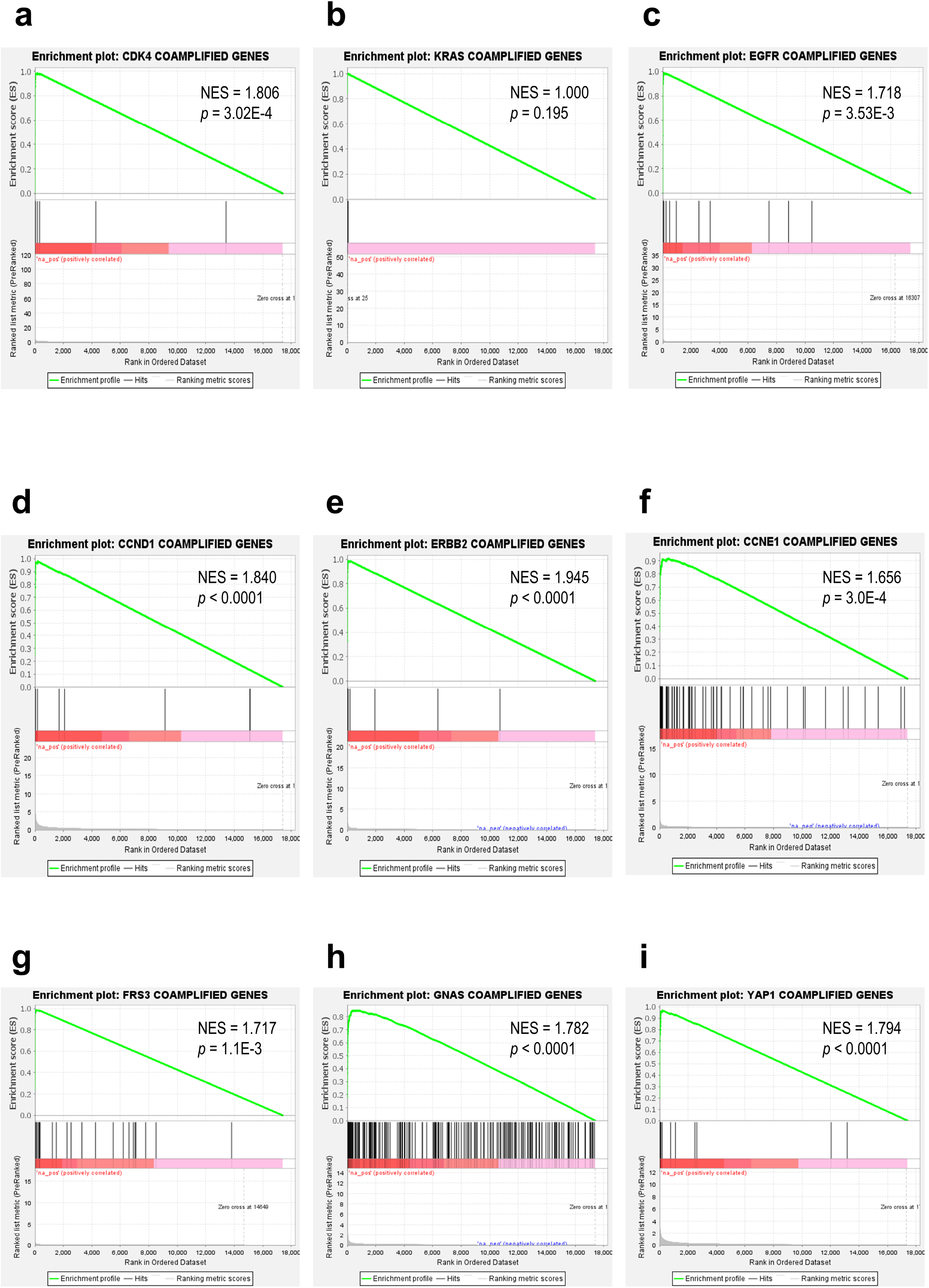
Preranked GSEA reveals an enrichment in coamplified genes among significant dependent genes for each gene amplification. Enrichment plots are shown for 9 of 16 amplifications analyzed: (**a**) *CDK4*, chr12q14 (**b**) *KRAS*, chr12p12 (**c**) *EGFR*, chr7p11 (**d**) *CCND1*, chr11q13 (**e**) *ERBB2*, chr17q11 (**f**) *CCNE1*, chr19q12 (**g**) *FRS3*, chr6p21 (**h**) *GNAS*, chr20q13 (**i**) *YAP1*, chr11q22. Normalized enrichment scores (NES) were used as a primary statistic to compare gene set enrichment results and the associated *p*-value were used to assess the statistical significance (*p* < 0.05).

**Supplementary Figure S5.**
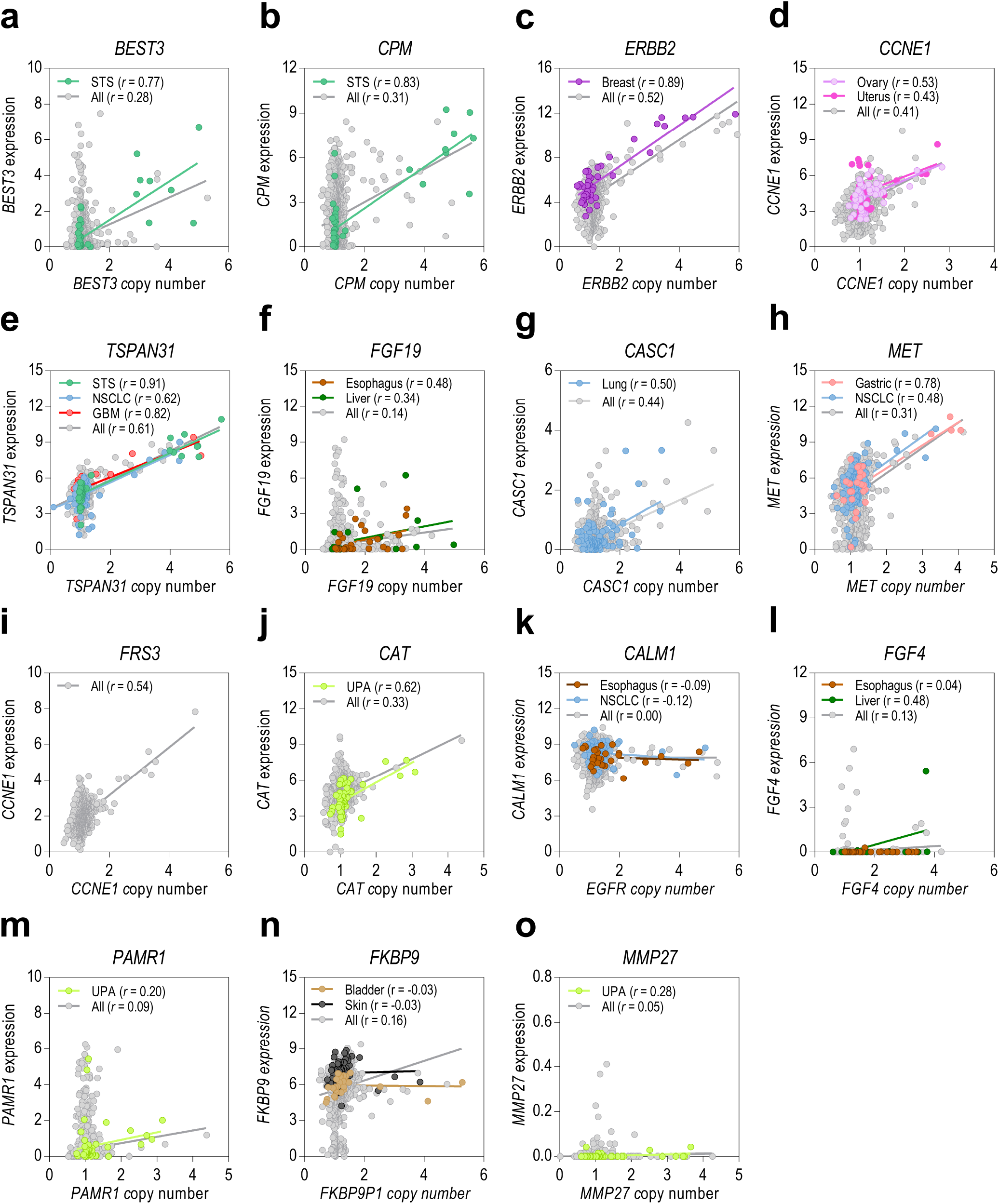
An increase in copy number does not always cause an increase gene expression in those prioritized genes. (**a-o**) Pearson’s correlations between gene relative copy number and relative gene expression are shown for those prioritized genes in cell lines from selected lineages. STS: soft-tissue sarcoma; NSCLC: non-small cell lung cancer; GBM: glioblastoma, UPA: upper aerodigestive tumors.

